# Gorlin syndrome-induced pluripotent stem cells form medulloblastoma with loss of heterozygosity in PTCH1

**DOI:** 10.1101/858555

**Authors:** Yu Ikemoto, Toshiyuki Miyashita, Michiyo Nasu, Hiromi Hatsuse, Kazuhiro Kajiwara, Katsunori Fujii, Toshino Motojima, Masashi Toyoda, Akihiro Umezawa

## Abstract

Gorlin syndrome is a rare autosomal dominant hereditary disease with high incidence of tumors such as basal cell carcinoma and medulloblastoma. Disease-specific induced pluripotent stem cells (iPSCs) have now been used as a model to analyze disease pathogenesis as well as an animal model. In this study, we generated iPSCs derived from fibroblasts of four patients with Gorlin syndrome (Gln-iPSCs) with a heterozygous mutation of the *PTCH1* gene. Gln-iPSCs from the four patients developed medulloblastoma in 100% (four out of four), a manifestation of Gorlin syndrome, in the teratomas after implantation into immunodeficient mice, but none (0/584) of the other iPSC-teratomas. One of the medulloblastomas had loss of heterozygosity in the *PTCH1* gene while benign teratoma, i.e. non-medulloblastoma part, did not, indicating a close clinical correlation between tumorigenesis in Gorlin syndrome patients and Gln-iPSCs.

## Introduction

Gorlin syndrome, a rare autosomal dominant disorder, is characterized by developmental defects in multiple organs or tissues such as skin (palmar or plantar pits), nervous system, eyes, endocrine systems and bones (bifid ribs). Gorlin syndrome is also associated with tumorigenesis such as development of basal cell carcinoma (BCC), medulloblastoma or keratocystic odontogenic tumor. Gorlin syndrome is caused by mutations in the *PTCH1* gene, a human homologue of *Drosophila patched*. PTCH1 is a member of hedgehog signaling components composed of hedgehog, SMO and GLI proteins. Hedgehog signaling regulates cell growth and development, and thus the disorder of this pathway gives rise to not only developmental anomalies but also diverse tumors such as those seen in Gorlin syndrome (1). Aberrant activation of hedgehog signaling causes basal cell carcinoma (2,3) and medulloblastoma (4,5).

Medulloblastoma occurs at an increased rate in mice with germline mutations in the *ptch1* gene. Although mice with *ptch1* deficiency are informative models for studies of Gorlin syndrome and medulloblastoma development, differences in underlying biology exist between humans and mice. Medulloblastoma forms in similar phenotype and anatomical location by haploinsufficiency of *ptch1* in mice, but do not exhibit loss of heterozygosity (6,7). Human disease-specific induced pluripotent stem cell (iPSC) has been used as one of human disease models to complement animal models (8,9). Basic schemes how to utilize patient iPSCs for disease mechanism studies have been well demonstrated (10,11). The power of iPSC technology for pathobiology studies is indeed revolutionary. Once established from any given patient, iPSCs serve as enduring resources for providing various functional cell types, eventually forever, which retain all the genomic information from the original patient. Using the system, we would be able to further investigate how disease-related phenotypes develop ‘in a dish’, or even test if novel therapeutic approaches can reverse the changes.

In this study, we focus on Gorlin syndrome, one of the well characterized disorders with mutations of Hedgehog signaling pathway. We have successfully generated medulloblastoma model with iPSCs derived from patients with Gorlin syndrome (Gln-iPSCs). Interestingly, Gln-iPSCs with a heterozygous germline mutation of *PTCH1* developed medulloblastoma with the secondary somatic mutation, i.e. LOH, in *PTCH1* in vivo. This iPSC model possibly serves pharmaceutical approach to find small molecules to treat medulloblastoma and Gorlin syndrome.

## Materials and Methods

### Ethical statement

Human cells in this study were obtained in full compliance with the Ethical Guidelines for Clinical Studies (Ministry of Health, Labor, and Welfare). The experimental procedure was approved by the Institutional Review Board (IRB) at National Center for Child Health and Development, Kitasato University and Chiba University Graduate School of Medicine.

### Human Cells

Patient cells were obtained from four patients diagnosed as Gorlin syndrome carrying a confirmed *PTCH1* mutation at the time of surgery (Table 1). Fibroblasts were grown from non-tumor tissues. Cells were cultured in 100-mm dishes (Becton Dickinson). All cultures were maintained at 37°C in a humidified atmosphere containing 95% air and 5% CO2. When the cultures reached subconfluence, the cells were harvested with a Trypsin-EDTA solution (cat# 23315, IBL CO., Ltd, Gunma, Japan), and re-plated at a density of 5 × 10^5^ cells in a 100-mm dish. Medium changes were carried out twice a week thereafter. Edom22-iPSCs (Edom22iPS#S31) (9) and human embryonic stem cells (SEES-5, SEES-6 and SEES-7) were used as controls for Gln-iPSCs (12).

**Table 1.** *PTCH1* mutations in the genomes from patients’ blood.

| Patient | Type of mutation | Nucleotide change | Amino acid change | Symptoms |
| --- | --- | --- | --- | --- |
| G11 | Frameshift | c.3130_3131dupGC | p.V1045LfsX23 | Macrocephaly, mental retardation, polydactyly of right lower extremity, palmar pits, rib anomaly |
| G12 | Frameshift | c.3130_3131dupGC | p.V1045LfsX23 | Palmar pits, odontogenic keratocysts of the jaw, multiple BCCs |
| G36 | Deletion of the whole *PTCH1* gene |  |  | Bifid ribs, kyphoscoliosis, macrocephaly, frontal bossing, hypertelorism |
| G72 | Frameshift | c.272delG<br>c.274delT*<sup>1</sup> | p.G91VfsX26<br>p.C92VfsX25 | Odontogenic keratocysts of the jaw, palmar pits, calcification of falx cerebri, stomach cancer |

\*1: Patient “G72” had a germline mutation, c.272delG, and a low prevalence of somatic mutation, c.274delT, that was derived from the allele with a germline mutation, but not from the wild-type allele (25).

### Generation of iPSCs

iPSCs were generated according to the method supplied with the CytoTune-iPS 2.0 Sendai Reprogramming Kit (MBL). Fibroblasts were seeded at 4.0×10^5^cells per well in a 6-well plate 24 h before infection. Sendai viruses expressing human transcription factors *OCT3/4, SOX2, KLF4*, and *c-MYC* were mixed in fibroblast medium to infect fibroblasts according to the manufacturer’s instructions. Six days after transfection, the transduced cells were detached using Trypsin/EDTA solution (Wako) and passaged onto irradiated mouse embryonic fibroblast feeder cells in fibroblast medium. On the next day, the medium was exchanged with human iPSC medium. Human iPSC medium contained KO-DMEM, KSR, GlutaMAX, NEAA, 2-Mercaptoethanol, Penicillin/Streptomycin, sodium pyruvate and bFGF (all from Invitrogen). From the next day, the medium was changed every day and the culture dishes were monitored for the emergence of iPSC colonies. When colonies were ready for transfer, they were picked up and expanded. Elimination of Sendai virus was confirmed by RT-PCR. Cells just after infection served a positive control. Sequences of the primers set are: forward primer, 5’-AGA CCC TAA GAG GAC GAA GA-3’; reverse primer, 5’-ACT CCC GSG GCG TAA CTC CGS AGT G-3’.

### iPSC culture

Human iPSCs were cultured onto a feeder layer of freshly plated gamma-irradiated mouse embryonic fibroblasts, isolated from ICR embryos at 12.5 gestations and passages 2 times before irradiation (30 Gy), in the iPSC media or in a feeder-free condition in StemFit AK02N (AJINOMOTO) in 6-cm dishes coated with 0.5 μg/cm^2^ vitronectin (Life Technologies). The cells were expanded using glass capillaries manually, or passaging by CTK solution (ReproCELL). When passaging the cells to feeder free, the cells were dissociated into single cells by treatment with using 0.5× TrypLE Select (Life Technologies) (1× TrypLE Select diluted 1:1 with 0.5 mM EDTA/PBS(-)) and replated at a density of 6cm dishes with StemFit media with 10 μM Y-27632 (Wako).

For EB formation, iPSC colonies were dissociated into single cells with accutase (Thermo Scientific, MA, USA) and then passaged into the low cell-adhesion 96 well plate dishes at a density of 10,000 cells/well in the iPSC medium without bFGF, and supplemented with ROCK inhibitor (12). After confirming EB formation on day 7, the EBs were harvested and passage to dishes coated with Basement Membrane Matrix (354234, BD Biosciences). Thereafter the EBs were maintained for 14 days and changed the iPSC medium without bFGF every other day.

### Quantitative RT-PCR

Total RNA was isolated from cells using the RNeasy Plus Mini Kit (QIAGEN). cDNA was synthesized from 1 mg of total RNA using Superscript III reverse transcriptase (Invitrogen) with random hexamers according to the manufacturer’s instructions. Template cDNA was PCR-amplified with gene-specific primer sets (Supplemental Table S1). RNA was extracted from cells using the RNeasy Plus Mini kit (QIAGEN). An aliquot of total RNA was reverse transcribed using an oligo (dT) primer. For the thermal cycle reactions, the cDNA template was amplified (ABI PRISM 7900HT Sequence Detection System) with gene-specific primer sets using the Platinum Quantitative PCR SuperMix-UDG with ROX (11743-100, Invitrogen) under the following reaction conditions: 40 cycles of PCR (95°C for 15 s and 60°C for 1 min) after an initial denaturation (95°C for 2 min). Fluorescence was monitored during every PCR cycle at the annealing step. The authenticity and size of the PCR products were confirmed using a melting curve analysis (using software provided by Applied Biosystems) and a gel analysis. mRNA levels were normalized using GAPDH as a housekeeping gene.

### Immunocytochemical analysis

Cells were fixed with 4% paraformaldehyde in PBS for 10 min at 4°C. After washing twice with PBS and treatment with 0.2% tritonX-100 in PBS for 10 min at 4°C, cells were pre-incubated with blocking buffer (Protein Block Serum Free solution, DAKO) for 30 min at room temperature, and then reacted with primary antibodies in 1% BSA in PBS for overnight at 4°C. Followed by washing with PBS, cells were incubated with secondary antibodies; anti-rabbit or anti-mouse IgG conjugated with Alexa 488 or 546 (15300) (Invitrogen) in 1% BSA in PBS for 1 h at room temperature. Then, the cells were counterstained with DAPI and mounted.

### Immunohistochemistry

Immunohistochemistry was performed as previously described (24). Paraffin sections were deparaffinized, dehydrated, and heated in Histofine Simple Stain MAX PO (MULTI) (Nichirei, Japan) for 20 min. After washing with distilled water, samples were placed in 1% hydrogen peroxide/methanol for 15 min to block endogenous peroxidase. Then, samples were incubated with blocking buffer (Protein Block Serum Free solution, DAKO) for 10 min at room temperature, The sections were then incubated at room temperature for 60 min in primary antibodies diluted with antibody diluent (Dako). The following primary antibodies against the antigens were used: Tuj-1 (1:300, Promega), Synaptophysin (1:400, DAKO), Nestin (1:200, Sigma-Aldrich), Ki-67 (1:100, Abcam), p53 (DAKO, 1:50). Then, they were washed three times with 0.01 M Tris buffered saline (TBS) solution (pH 7.4) and incubated with goat anti-mouse or anti-rabbit immunoglobulin labeled with dextran molecules and horseradish peroxidase (EnVision, Dako) at room temperature for 30 min. After washing with TBS, they were incubated in 3,3’-diaminobenzidin in substrate-chromogen solution (Dako) for 5-10 min. Negative controls were performed by omitting the primary antibody. The sections were counterstained with hematoxylin.

### Karyotypic analysis

Karyotypic analysis was contracted out at Nihon Gene Research Laboratories Inc. (Sendai, Japan). Metaphase spreads were prepared from cells treated with 100 ng/mL of Colcemid (Karyo Max, Gibco Co. BRL) for 6 h. The cells were fixed with methanol: glacial acetic acid (2:5) three times, and dropped onto glass slides (Nihon Gene Research Laboratories Inc.). Chromosome spreads were Giemsa banded and photographed. A minimum of 10 metaphase spreads were analyzed for each sample, and karyotyped using a chromosome imaging analyzer system (Applied Spectral Imaging, Carlsbad, CA).

### Short tandem repeat analysis

Short tandem repeat analysis was contracted out at BEX Inc. (Tokyo,Japan) and used of the PowerPlex® 16 System(Promega). One primer specific for D3S1358, TH01, D21S11, D18S51, and Penta E is labeled with fluorescein (FL); one primer specific for D5S818, D13S317, D7S820, D16S539, CSF1PO, and Penta D is labeled with 6-carboxy-4′,5′-dichloro-2′,7′-dimethoxy-fluorescein (JOE); and one primer specific for Amelogenin, vWA, D8S1179, TPOX, and FGA is labeled with carboxy-tetramethylrhodamine (TMR). Genotyping of cell lines is analyzed by co-amplification of all sixteen loci and three-color detection.

### Teratoma formation

Gln-iPSCs were harvested by accutase treatment, collected into tubes, and centrifuged. The same volume of Basement Membrane Matrix (354234, BD Biosciences) was added to the cell suspension. The cells (>5.0×10^6^) were subcutaneously inoculated into immunodeficient mice (BALB/cAJcl-nu/nu, CREA, Tokyo, Japan). After 6 to 12 weeks, the resulting tumors were dissected and fixed with formalin. Paraffin-embedded tissue was sliced and stained with hematoxylin and eosin (HE). The operation protocols were accepted by the Laboratory Animal Care and the Use Committee of the National Research Institute for Child and Health Development, Tokyo.

### Laser Microdissection

Paraffin sections were cut as 10-micrometer sections onto PEN-membrane slides (Leica Microsystems, Wetzlar, Germany). Tissue sections were stained with hematoxylin. After dry out the tissue, target area was extracted by use of a Leica LMD6500 (Leica).

### Mutational analysis

DNA was extracted using a DNeasy Blood & Tissue Kit (QIAGEN), QIAamp DNA mini kit (QIAGEN) or QIAamp DNA blood midi kit (QIAGEN). Genomic DNA was PCR-amplified with specific primer sets for the *PTCH1* gene (Supplemental Table S2). Amplified products were gel-purified using a QIAEX II gel extraction kit (QIAGEN) and cycle sequenced with a BigDye Terminator v3.1 Cycle Sequencing Kit (Applied Biosystems) in both directions. The sequence was analyzed on a 3130 Genetic Analyzer (Applied Biosystems). For some analyses, PCR products were subcloned into the pGEM-T Easy vector (Promega) and the inserts were sequenced.

## Results

### Generation and characterization of Gln-iPSCs

We generated iPSCs from human cells with a mutation in the *PTCH1* gene by Sendai virus infection-mediated expression of *OCT4/3, SOX2, KLF4*, and *c-MYC* (Figure 1A). When the reprogramming factors OCT4/3, SOX2, KLF4 and c-MYC were introduced into 4.0 × 10^5^ cells, iPSCs generated from four patients with Gorlin syndrome were successfully generated and designated as G11-, G12-, G36- and G72-iPSC. Efficiency of iPSC generation was then calculated as “iPSC colonies generated/fibroblasts exposed to virus”. The efficiency of the iPSC colony generation was relatively high, i.e. 0.1% to 1.0%, compared with that of iPSCs (Edom22-iPSCs) generated from healthy individuals. Morphological characteristics of Gln-iPSCs, *i*.*e*. flat and aggregated colonies, were similar to those of other intact iPSCs and ESCs (Figure 1B). RT-PCR analysis revealed elimination of the Sendai virus (Figure 1C). Immunocytochemical analyses demonstrated expression of the pluripotency-associated markers, *i*.*e*. SSEA-4, TRA-1-60, SOX2, NANOG, and OCT4/3, which was consistent with the profile observed in hESCs (Figure 1D). The expression profiles of stem cell-associated genes were examined with qualitative RT-PCR to confirm the iPSC-characters. Expressions of pluripotency-associated genes, such as *SOX2, OCT4/3, DNMT3B, NANOG*, and *TERT*, were detected in all Gln-iPSC clones to a similar extent of those in control human embryonic stem cells (12) and healthy donor-derived iPSCs (13) (Figure 1E). To evaluate whether Gln-iPSCs maintained their pluripotency in vitro, we performed EB assays. EBs differentiated from the cells of SEES-4, SEES-5, SEES-6, and SEES-7 cells expressed markers associated with the three major germ layers: TUJ1 (ectoderm), αSMA (mesoderm), and AFP (endoderm) (Figure 1F). Short tandem repeat (STR) analysis showed clonality between the respective iPSC lines and their parental cells (Table 2). Gln-iPSCs cells showed intact karyotypes (Figure 1G).

**Table 2.** STR analyses of Gorlin iPSCs.

| Locus | G11 fibroblasts |  | G11 iPSC |  | G12 fibroblasts |  | G12 iPSC |  | G36 fibroblasts |  | G36 iPSC |  | G72 fibroblasts |  | G72 iPSC |  |
| --- | --- | --- | --- | --- | --- | --- | --- | --- | --- | --- | --- | --- | --- | --- | --- | --- |
| D3S1358 | 15 | 16 | 15 | 16 | 15 | 16 | 15 | 16 | 15 |  | 15 |  | 15 | 16 | 15 | 16 |
| TH01 | 9 |  | 9 |  | 6 | 9 | 6 | 9 | 9 |  | 9 |  | 9 |  | 9 |  |
| D21S11 | 31.2 |  | 31.2 |  | 30 | 31.2 | 30 | 31.2 | 29 | 30 | 29 | 30 | 30 |  | 30 |  |
| D18S51 | 15 |  | 15 |  | 15 |  | 15 |  | 14 | 18 | 14 | 18 | 15 | 16 | 15 | 16 |
| Penta_E | 14 | 17 | 14 | 17 | 14 | 20.2 | 14 | 20.2 | 11 | 17 | 11 | 17 | 8 | 14 | 8 | 14 |
| D5S818 | 10 | 11 | 10 | 11 | 9 | 11 | 9 | 11 | 10 | 11 | 10 | 11 | 10 | 11 | 10 | 11 |
| D13S317 | 8 | 9 | 8 | 9 | 8 |  | 8 |  | 8 | 9 | 8 | 9 | 8 | 11 | 8 | 11 |
| D7S820 | 10 | 11 | 10 | 11 | 10 | 12 | 10 | 12 | 11 |  | 11 |  | 11 |  | 11 |  |
| D16S539 | 10 | 11 | 10 | 11 | 11 | 13 | 11 | 13 | 10 | 12 | 10 | 12 | 9 | 12 | 9 | 12 |
| CSF1PO | 11 | 12 | 11 | 12 | 11 | 12 | 11 | 12 | 11 | 12 | 11 | 12 | 9 | 11 | 9 | 11 |
| Penta_D | 10 |  | 10 |  | 9 | 10 | 9 | 10 | 9 | 13 | 9 | 13 | 9 | 10 | 9 | 10 |
| AMEL | X | Y | X | Y | X |  | X |  | X |  | X |  | X |  | X |  |
| vWA | 16 | 18 | 16 | 18 | 16 | 18 | 16 | 18 | 16 |  | 16 |  | 13 | 19 | 13 | 19 |
| D8S1179 | 13 |  | 13 |  | 13 | 14 | 13 | 14 | 10 | 11 | 10 | 11 | 11 | 12 | 11 | 12 |
| TPOX | 8 | 11 | 8 | 11 | 11 |  | 11 |  | 8 | 11 | 8 | 11 | 8 |  | 8 |  |
| FGA | 22 | 24 | 22 | 24 | 22 | 24 | 22 | 24 | 21 | 23 | 21 | 23 | 23 | 25.2 | 23 | 25.2 |

**Figure 1.**
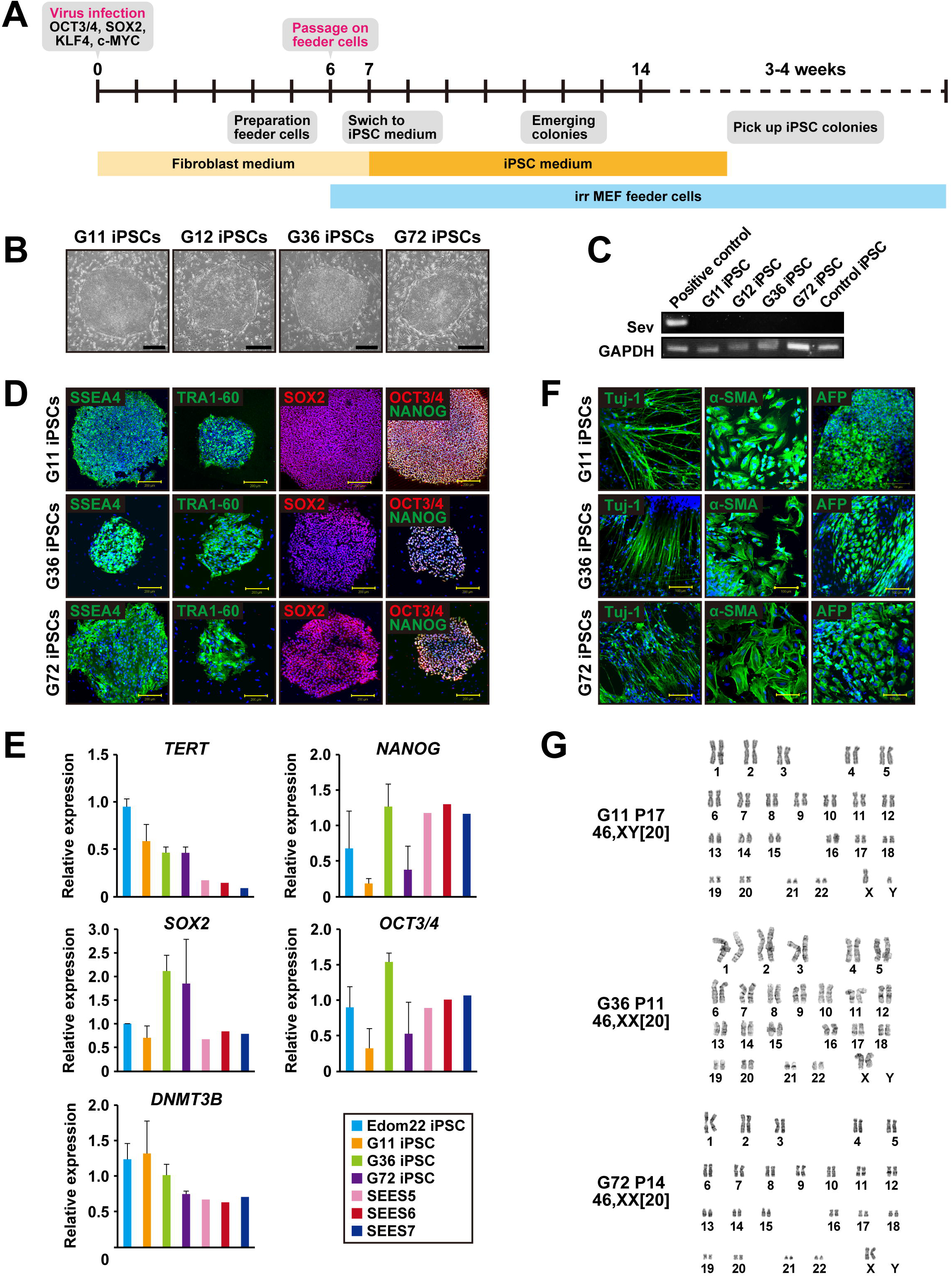
Generation of iPSCs from fibroblasts of the patients with Gorlin syndrome. A. Protocol for iPSC generation. B. Phase-contrast microphotographs of Gln-iPSCs (G11, G12, G36, G72). C. RT-PCR analysis of the Sendai virus. D. Immunocytochemical analysis of Gln-iPSCs using antibodies to NANOG, OCT4/3, SOX2, SSEA4, and TRA1-60. E. Expression of the endogenous *TERT, NANOG, SOX2, OCT4/3*, and *DNMT3B* genes. F. in vitro differentiation of Gln-iPSCs into three germ layers. Immunocytochemical analysis of Gln-iPSCs using antibodies to Tuj-1, α-smooth muscle actin (SMA) and α-fetoprotein (AFP). Expression of the endogenous *TERT, NANOG, SOX2, OCT4/3*, and *DNMT3B* genes. G. Karyotypes of Gln-iPSCs at the indicated passage number. Numbers in brackets indicate the number of cells analyzed.

### Sequencing analysis of the *PTCH1* gene

*PTCH1* mutations of iPSC lines established from different 4 patients were determined by the direct sequencing method (Figure 2A, Table 1). G12-iPSCs and G11-iPSCs were derived from cells of mother and son, respectively, and had the same heterozygous mutation, c.3130_3131dupGC, in exon 18, resulting in the frameshift. G36-iPSCs had a heterozygous deletion of the whole *PTCH1* gene. G72-iPSCs had heterozygous mutation, c.272G, in exon 2, resulting in the frameshift. All mutations identified in iPSCs were identical to those in their parental fibroblasts.

**Figure 2.**
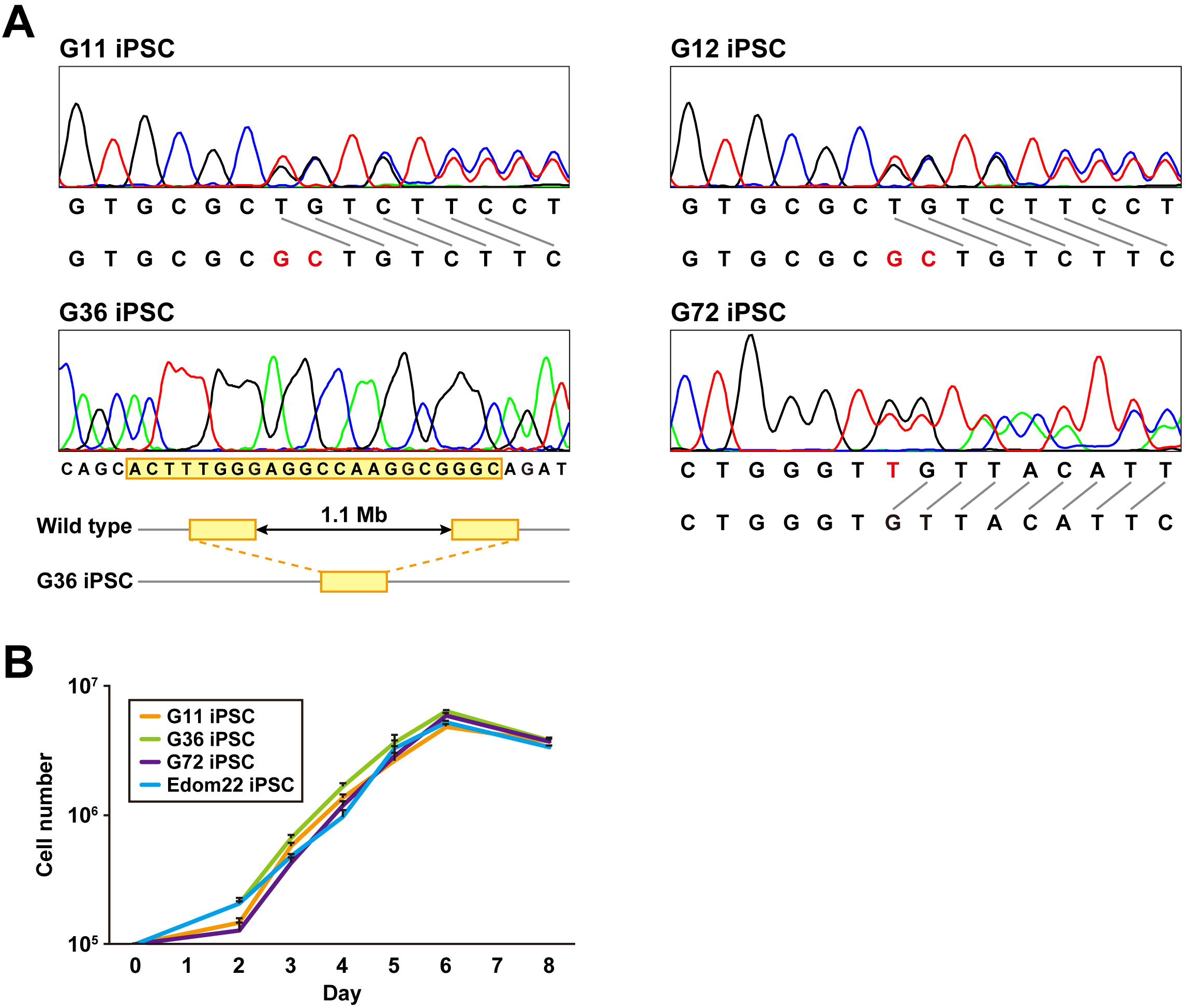
Genomic analysis and cell proliferation assay of Gln-iPSCs. A. Sequencing analysis of the *PTCH1* gene in Gln-iPSCs. B. Growth curves of Gln-iPSCs. Cell numbers of Gln-iPSCs (G11, G36, G72) and Edom22-iPSCs (control iPSCs) were counted at the indicated days after cells (1.0 × 10^5^ cells/dish) were seeded on vitronectin-coated 6-well plates.

### Characterization of Gln-iPSCs

The proliferative capacity of three Gln-iPSC clones (G11, G36, G72) was measured and compared with that of Edom22-iPSCs (Figure 2B). No significant differences in proliferation rates were detected between the Gln-iPSC clones and Edom22-iPSCs.

### Teratoma formation

To address whether the Gln-iPSCs have the competence to differentiate into specific tissues, teratoma formation was tested by implantation of Gln-iPSCs in the subcutaneous tissue of immunodeficient Balb/c nu/nu mice. Gln-iPSCs produced teratomas within 6-12 weeks after the implantation. Histological analysis of paraffin-embedded sections demonstrated that the three primary germ layers were generated as shown by the presence of ectodermal, mesodermal, and endodermal tissues in the teratoma (Figure 3A), implying that Gln-iPSCs have potential for multilineage differentiation in vivo. Area of neuroepithelium, retina, and retinal pigmented epithelium was relatively large, compared with that of other mesodermal and endodermal components.

**Figure 3.**
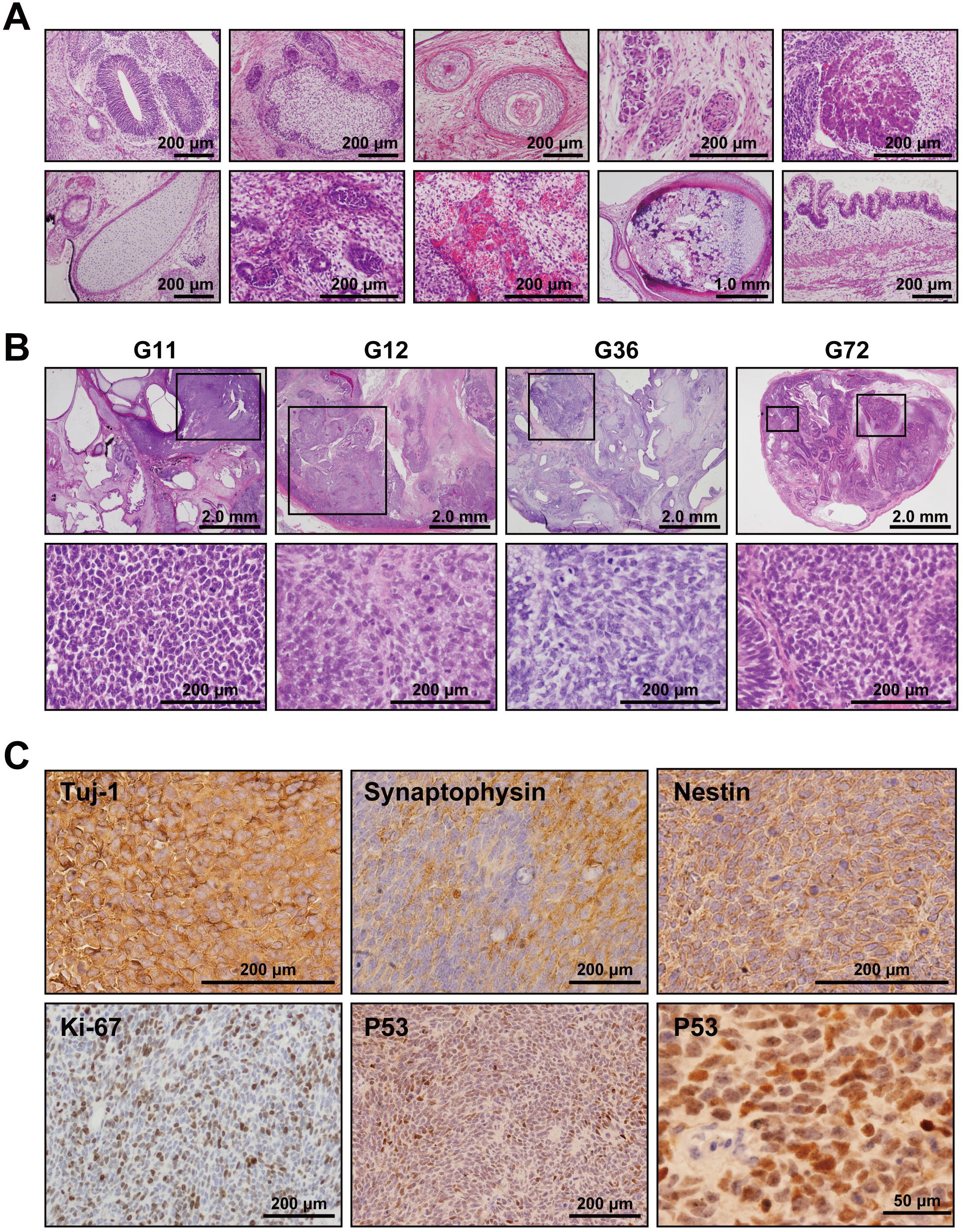
Medulloblastoma in Gln iPSC-teratoma. A. Histology of teratoma generated by Gln-iPSCs. Upper panels from left to right: ectodermal glia and neuroepithelium; epidermis with hair follicles; epidermis with keratinization; ganglia; hepatocytes. Lower panels from left to right: cartilage; glomerulus-like structure; capillary vessels; bone and cartilage; intestinal epithelium. B. Medulloblastomas were generated in the teratomas by Gln-iPSCs (G11, G12, G36, G72). Upper panels: low power view of the teratomas. Lower panels: high power view of medulloblastoma parts. Medulloblastomas were shown in the squares of the upper panels. C. Immunohistochemical analysis of medulloblastoma using antibodies to Tuj-1, synaptophysin, nestin, Ki-67, and p53.

### Medulloblastoma formation

Medulloblastoma was formed in all the teratoma generated by all Gln-iPSC clones from 4 different patients (Figure 3B). Diagnosis of medulloblastoma was confirmed by the certified pathologists of the two independent organizations. The tumor cells stained strongly positive for TUJ1, Synaptophysin, NESTIN, Ki67 and p53. Sequencing analysis of the microdissected cartilage and medulloblastoma revealed heterozygous mutation of the *PTCH1* gene and LOH of the *PTCH1* gene, respectively (Figure 4).

**Figure 4.**
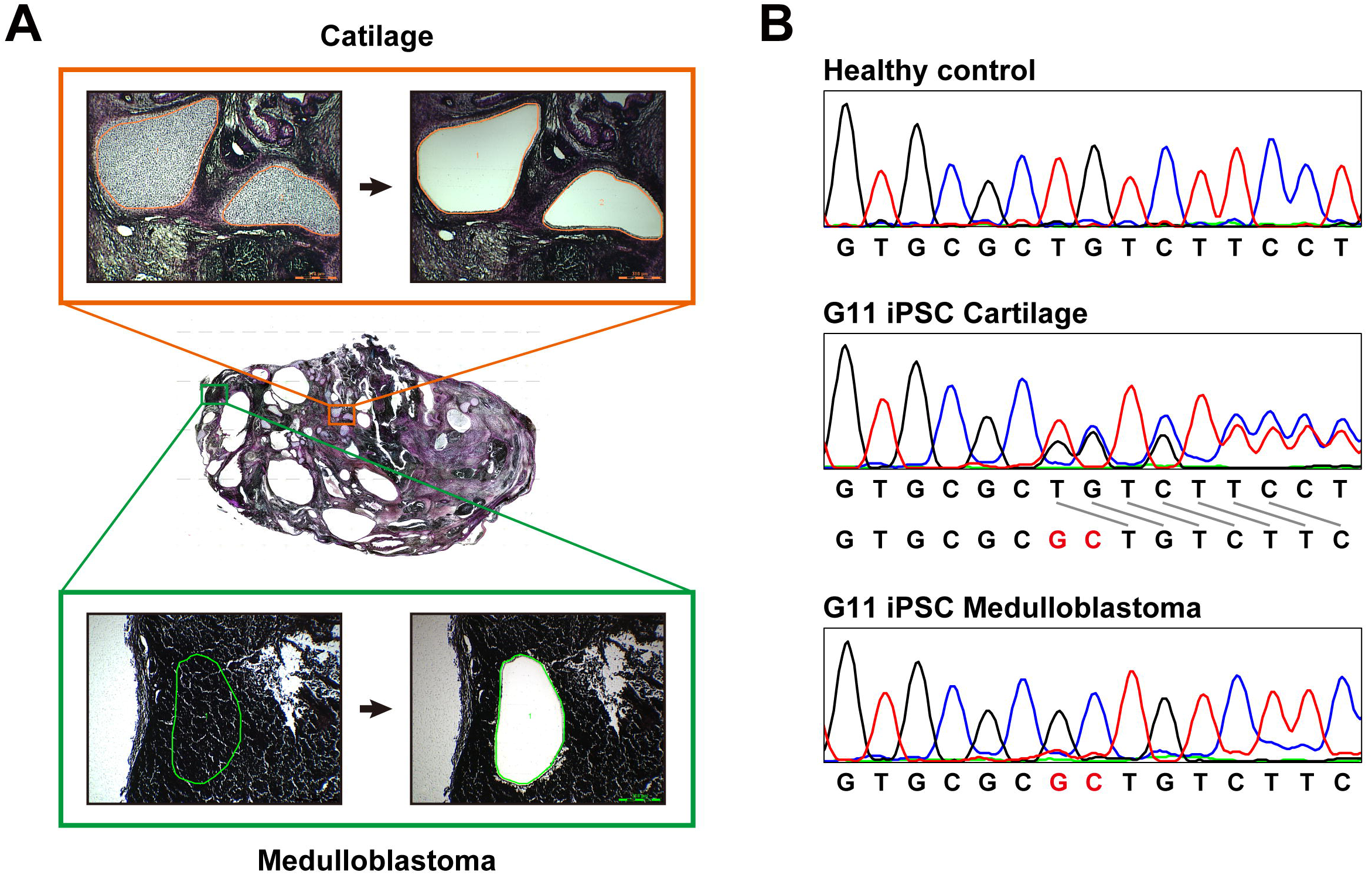
Sequencing analysis of the *PTCH1* gene in medulloblastoma. A. Microdissection of medulloblastoma and cartilage. Genomic DNA was isolated from the microdissected medulloblastoma, and applied to direct sequencing analysis. Genomic DNA was also isolated from the cartilage for comparison. B. Direct sequencing analysis of genomic DNA from unaffected donor (control), G11-iPSC cartilage and G11-iPSC medulloblastoma. G11-iPSC cartilage had a duplication of “GC” (shown in red), and G11-iPSC medulloblastoma had LOH of the *PTCH1* gene.

## Discussion

Human pluripotent stem cells deficient for a gene can be generated in two ways: Disruption of the gene in human ESCs or intact iPSCs by genetic manipulation with bacterial artificial chromosome and derivation of disease-specific iPSCs from patients with germline mutations. Patient iPSCs serve as disease model cells for clarification of pathogenic mechanisms and for screening novel compounds to treat the disease. In this study, we generated iPSCs from fibroblasts of a human Gorlin syndrome patient. Despite the comparable proliferation activity, Gln-iPSCs consistently developed medulloblastoma, i.e. Gorlin syndrome-associated tumor, with the secondary somatic mutation of the *PTCH1* gene.

### Modeling cancer with iPSCs from patients with germline mutation

Disease iPSCs have been analysed in context of tumorigenesis in Li-Fraumeni syndrome and hereditary breast-ovarian cancer syndrome (14,15). These two cancer-prone genetic disorder-derived iPSCs exhibit disease-specific phenotypes, but do not develop cancers in vivo. High frequency of medulloblastoma generation in Gorlin syndrome iPSC-derived teratomas was against of our unexpectation because secondary somatic mutation was detected from the specimens of the four patients with the independent germline mutations, implying that Gorlin syndrome iPSCs have genomic stability that is associated with LOH of the *PTCH1* gene albeit lack of chromosomal aberration even after long-term cultivation of Gln-iPSCs. The secondary somatic mutation might occur before implantation, however the percentage of Gln-iPSCs with LOH of the *PTCH1* gene failed to be detected. Area of medulloblastoma was not limited in the teratomas, suggesting in vivo growth advantage of medulloblastoma generated by Gln-iPSC with LOH of the *PTCH1* gene.

Formation of BCC in Gln-iPSC-derived teratoma is also expected because BCC of both sporadic and familial BCC is correlated with LOH of the *PTCH1* gene as is the cases of medulloblastoma (16). Patients with Gorlin syndrome clinically have medulloblastoma at children, i.e. between the ages of 0 and 4 years. In contrast, they start to have BCC after age 20 in Japan and probability of developing BCC increases with age (17). The frequency of BCC in Japanese patients of over 20 years exceeds 50%. This late onset of BCC is possibly related with lack of BCC in the Gln-iPSC teratoma. Additional secondary mutation except for *PTCH1* LOH could be required for generation of BCC. The Gln-iPSC teratoma formed in the subcutaneous tissue of the experimental mice in the animal facility with fluorescent lighting and was therefore unexposed to chemical mutagen and sunlight/irradiation. Such special circumstance of the Gln-iPSC teratoma may be attributed to lack of BCC.

### Animal models of Gorlin syndrome

As the most common malignant pediatric brain tumor, murine models were generated. Ptc-/- murine model (B6D2F1 background) develop medulloblastoma at 14% (4). Through experiments by crossing the model mice with p53-/- mice, genomic instability possibly contributes to medulloblastoma generation. The high incidence of medulloblastoma increases availability of the model for preclinical studies (18). Medulloblastoma in mice with haploinsufficiency of *ptch1* forms in similar phenotype and anatomical location of human medulloblastoma, and therefore mice genetically engineered to have *ptch1* deficiency are informative models for studies of Gorlin syndrome and Medulloblastoma development. However, medulloblastoma in ptc+/- mice do not exhibit LOH unlike Gln-iPSC-induced medulloblastoma. The differences between murine individual models and human Gln-iPSC models are unclear, and could be attributed to the differences between mouse and human or between the individual and cellular levels.

### Acceleration of HH signaling in correlation with iPSC generation

The warranted quality of Gln-iPSCs indicates that growth and differentiation capability of iPSCs remains unaffected albeit accelerated hedgehog signaling. Relatively high efficiency of Gln-iPSC colony generation possibly relates accelerated reprogramming of human somatic cells to iPSCs by sonic hedgehog (13,19,20). Participation of sonic hedgehog signaling in neural differentiation of human pluripotent stem cells may also be related to enlarged area of neuroectodermal component in the Gln-iPSC teratoma and medulloblastoma formation (21–23).

Gln-iPSCs can be a good model of Gorlin syndrome to identify biomarkers, and thereby used for drug screening to prevent tumors for medulloblastoma and BCC. Consistent generation of medulloblastoma in Gln-iPSC teratoma is also available for drug screening of anticancer drug. Moreover, Gln-iPSCs may serve a model to elucidate mechanism for LOH of the *PTCH1* gene in patients with Gorlin syndrome.

## Acknowledgements

We would like to express our sincere thanks to K. Miyado and H. Akutsu for fruitful discussion, to M. Ichinose for providing expert technical assistance, to C. Ketcham for English editing and proofreading, and to E. Suzuki and K. Saito for secretarial work.

