## Supplemental Table 1 for "Gorlin syndrome-induced pluripotent stem cells form medulloblastoma with loss of heterozygosity in PTCH1"

**Supplemental Table S1. Primer sets for the pluripotency-associated genes.**

| | Forward (5'→3') | Reverse (5'→3') |
|---|---|---|
| *OCT4/3* | TGTACTCCTCGGTCCCTTTC | TCCAGGTTTTCTTTCCCTAGC |
| *NANOG* | CAGTCTGGACACTGGCTGAA | CTCGCTGATTAGGCTCCAAC |
| *SOX2* | ATGGGTTCGGTGGTCAAGT | GGAGGAAGAGGTAACCACAGG |
| *DNMT3B* | GGAAATTAGAATCAAGGAAATACGA | AATTTGTCTTGAGGCGCTTG |
| *TERT* | GGAGCAAGTTGCAAAGCATTG | TCCCACGACGTAGTCCATGTT |
| *GAPDH* | TGTTGCCATCAATGACCCCTT | CTCCACGACGTACTCAGCG |
