## Supplemental Table 2 for "Gorlin syndrome-induced pluripotent stem cells form medulloblastoma with loss of heterozygosity in PTCH1"

**Supplemental Table S2. Primer sets to detect mutations of the PTCH1 gene.**

| | Forward (5'→3’) | Reverse (5'→3’) |
|---|---|---|
| G11(G12) | AACTGTGATGCTCTTCTACCCTGG | TCTTTCTGCAGCCGGGAAGTTTT |
| G36 | CAACACCCAATTCTGGATAC | AAATCAGAGCCTGCATTCGC |
| G72 | CACTCCTCCCTTCTGCTTCG | TCTGCCACGTATCTGCTCAC |
